# Nitrite lowers the oxygen cost of ATP supply in skeletal muscle cells by stimulating the rate of glycolytic ATP synthesis

**DOI:** 10.1101/2022.03.30.486447

**Authors:** Anthony G. Wynne, Charles Affourtit

## Abstract

Dietary nitrate lowers the oxygen cost of human exercise. This effect has been suggested to result from stimulation of coupling efficiency of skeletal muscle oxidative phosphorylation by reduced nitrate derivatives. In this paper, we report the acute effects of sodium nitrite on the bioenergetic behaviour of L6 myocytes. At odds with improved efficiency of mitochondrial ATP synthesis, extracellular flux analysis reveals that a ½-hour exposure to NaNO_2_ (0.1 – 5 µM) significantly decreases mitochondrial coupling efficiency in static myoblasts and tends to lower it in spontaneously contracting myotubes. Unexpectedly, NaNO_2_ stimulates the rate of glycolytic ATP production in both myoblasts and myotubes. Increased ATP supply through glycolysis does not emerge at the expense of oxidative phosphorylation, which means that NaNO_2_ acutely increases the rate of overall myocellular ATP synthesis, highly significantly so in myoblasts and tending towards significance in contractile myotubes. Notably, NaNO_2_ exposure shifts myocytes to a more glycolytic phenotype. Mitochondrial oxygen consumption does not decrease after NaNO_2_ exposure, and non-mitochondrial respiration tends to drop. When total ATP synthesis rates are normalised to total *cellular* oxygen consumption rates, it thus transpires that NaNO_2_ lowers the oxygen cost of ATP supply in L6 myocytes.

## Introduction

Inorganic nitrate 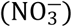 is found in beetroot and leafy vegetables such as spinach and rocket [1] and protects against cardiovascular disease [2]. This protection is afforded by nitric oxide formed through reduction of the dietary nitrate [3]. After ingestion, nitrate is first reduced to nitrite by nitrate reductases in bacteria that occupy the posterior part of the tongue [4]. Salivary nitrite is taken up into the circulation lifting the plasma nitrite level up to 600 nM [5]. At this relatively high concentration, and at a low pH and oxygen tension, nitrite can then be reduced to nitric oxide, possibly catalysed by xanthine oxidase [6] or deoxyhaemoglobin [7].

Dietary nitrate has been found to lower the oxygen cost of exercise, as it decreases the respiratory activity required for a set rate of skeletal muscle work [8,9]. This interesting finding challenges exercise physiology dogma that oxygen uptake for any individual is fixed at a set workload irrespective of age, fitness, diet or pharmacological intervention [10], and has provoked much new research [11]. Meta-analysis of this research confirms that nitrate enhances muscle performance in different exercise contexts [12,13], but highlights the need for mechanistic understanding. Such understanding may explain why the nitrate benefit does not extend to all types of exercise, why benefit is limited to certain human groups, and why any tested group contains distinct responders and non-responders [14].

The mechanism by which dietary nitrate improves muscle performance remains incompletely understood, but many models predict increased efficiency of myocellular bioenergetics [14]. A lowered oxygen cost of exercise has been attributed to raised efficiency of oxidative ATP supply [15], but this notion is controversial [16,17]. Dietary nitrate also improves contractile properties of human skeletal muscle [18], which suggests it lowers the ATP cost of work. These mechanisms are not mutually exclusive, as efficiency of oxidative ATP production and ATP consumption may both be increased [19]. It appears widely accepted that the exercise effects of dietary nitrate are mediated by nitric oxide [11], but the evidence is circumstantial. As nitrate/nitrite exposure increases the intracellular level of nitrate, nitrite and nitric oxide in human myocytes [20], any of these species may contribute, in principle, to enhanced muscle function.

Reasoning that nitrate may not be reduced to nitric oxide under the conditions that prevail during the dietary supplementation, and indeed during the submaximal intensity exercise it benefits most, we set out to establish how nitrite, i.e., the reduction intermediate between nitrate and nitric oxide, affects the bioenergetic behaviour of cultured skeletal muscle cells. Quantifying real-time ATP supply rates by extracellular flux analysis [21], we report here that sodium nitrite acutely stimulates the rate of glycolytic ATP production in rat L6 myoblasts as well as in spontaneously contracting L6 myotubes. Since this stimulation does not occur at the expense of ATP synthesis through oxidative phosphorylation, the oxygen cost of total ATP supply is lowered by nitrite.

## Material and methods

### Tissue culture

Clonal rat myoblasts (L6.C11) were obtained from the European Collection of Cell Culture. These L6 myoblasts were seeded 24 h before experiments at 30,000 cells per well on either XF24 tissue culture plates (Agilent-Seahorse) for extracellular flux analysis, or on 96-well plates (Corning) for glucose uptake assays. Cells were studied between passages 11 and 20 and were grown in DMEM supplemented with 5 mM glucose, 10% (v/v) fetal bovine serum under a humidified carbogen atmosphere of 5% CO_2_ and 95% air at 37 ºC. To allow cell differentiation, myoblasts were cultured to complete confluence in fully supplemented DMEM on XF24 or 96-well plates, at which point the serum concentration was decreased to 2% (v/v). When cells were cultured in this ‘light’-serum medium for 14 days – i.e., longer than the 8-10 days we stuck to before [22] – and growth medium was refreshed every 2-3 days, the L6 myoblasts differentiated into myotubes that contract spontaneously.

### Cellular bioenergetics

Oxygen consumption and medium acidification by myoblasts and spontaneously contracting myotubes were measured with a Seahorse XF24 extracellular flux analyser (Agilent). Before the assay, cells were washed into a Hepes-buffered Krebs-Ringer medium (KRH) comprised of 135 mM NaCl, 3.6 mM KCl, 2 mM Hepes (pH 7.4), 0.5 mM MgCl_2_, 1.5 mM CaCl_2_, 0.5 mM NaH_2_PO_4_, and 5 mM glucose. Cells were allowed to equilibrate in this medium for 30 min under air at 37 ºC, and were then transferred to the XF24 analyser for another 12-min equilibration. Oxygen consumption (OCR) and extracellular acidification (ECAR) rates were recorded 3x (measurement cycle: 2-min mix, 2-min wait and 3-min measure) at which point 0.1 – 5 µM NaNO_2_ was added; KRH and NaNO_3_ (up to 500 µM) were used as controls. Following about ½-h (26 min) incubation, 5 µg/mL oligomycin, 1 µM carbonyl cyanide-p-trifluoromethoxyphenylhydrazone (FCCP) or N5,N6-bis(2-fluorophenyl)[1,2,5]oxadiazolo[3,4-b]pyrazine-5,6-diamine (BAM15), and 1 µM rotenone with 1 µM antimycin A were added sequentially to inhibit the ATP synthase, uncouple oxidative phosphorylation, and inhibit the mitochondrial electron transfer chain, respectively. OCRs (calculated from time-resolved oxygen concentrations applying the *Akos* algorithm [23]) and ECARs were normalised to the number of nuclei quantified by 4’,6-diamidino-2-phenylindole (DAPI) staining as described before [24]. It was thus determined that each well contained on average 27K myoblast nuclei – i.e., within experimental variation of the 30K/well seeding density – or 39K myotube nuclei. OCRs resistant to rotenone and antimycin A were subtracted from all other OCRs to correct for non-mitochondrial oxygen uptake. Mitochondrial respiration coupled to ATP synthesis or associated with proton leak was gauged from the oligomycin-sensitive and oligomycin-resistant OCRs, respectively, while mitochondrial respiratory capacity was estimated from uncoupled respiration. Coupling efficiency of oxidative phosphorylation was defined as the percentage of basal respiration used to make ATP, and cell respiratory control as the ratio between uncoupled and oligomycin-inhibited respiration [25].

The rates of glycolytic and mitochondrial ATP synthesis were derived from cellular oxygen uptake and medium acidification as described by Mookerjee *et al*. [21], assuming myocyte energy metabolism was fuelled exclusively by glucose. The mitochondrial ATP supply rate was calculated from mitochondrial respiration coupled to ATP synthesis using a P/O ratio for glucose-fuelled TCA cycle turnover of 0.12 and a P/O ratio for glucose-fuelled oxidative phosphorylation of 2.50 [21], thus assuming that glycolytic reducing power is transferred to mitochondria by the malate-aspartate shuttle alone. ATP-synthesis-coupled respiration was approximated from the oligomycin-sensitive oxygen consumption after correcting for the small underestimation owing to oligomycin-induced hyperpolarisation of the mitochondrial inner membrane [21]. Glycolytic ATP synthesis was defined as the net ATP produced during glucose breakdown to pyruvate irrespective of pyruvate’s destiny. The glycolytic ATP supply rate was thus calculated from medium acidification to account for pyruvate that was reduced to lactate^−^ and H^+^ (ATP:lactate = 1:1) and from mitochondrial respiration to account for the pyruvate that was oxidised to bicarbonate (ATP:O_2_ = 0.33:1). A KRH buffering power of 0.681 mpH x µM^-1^ H^+^ was used to convert the ECAR to a proton production rate (PPR) assuming an effective XF24 measurement volume of 22.7 µL (25). The PPR was corrected for medium acidification accounted for by bicarbonate^−^ plus H^+^ that emerges from pyruvate oxidation to reflect contribution of lactate^−^ and H^+^ only [26]. Lactate production rates were also measured directly by lactate dehydrogenase assay of medium samples taken after completion of the XF run, and these rates were normalised to total protein [26].

### Glucose uptake

2-Deoxyglucose uptake by myoblasts and myotubes was measured as described previously [22]. Cells were assayed and washed in glucose-free KRH (see previous section), and were lysed in buffer containing 0.1 N HCl and 0.1% (w/v) Triton X100 at room temperature. This acidic lysis method offered a higher signal-to-noise ratio than the high-temperature alkaline procedure we used before [22].

### Statistical analysis

Differences between myocellular differentiation state and bioenenergetic condition as well as the NaNO_2_ and NaNO_3_ effects were evaluated for statistical significance using GraphPad Prism (version 9 for Windows), applying tests that are specified in the figure legends, and using an ordinary least square method for the linear regression analysis.

## Results

### Contractile myotubes rely more strongly on glycolytic ATP supply than static myoblasts

Prolonged growth of L6 myocytes in low-serum medium leads to formation of spontaneously contracting myotubes (https://www.youtube.com/watch?v=w5UU5UiEi3Q). We subjected these contractile cells to extracellular flux analysis to compare their bioenergetic behaviour with that of resting myoblasts. In both differentiation states, the basal OCR decreases when mitochondrial ATP synthesis is inhibited by oligomycin. Mitochondrial uncoupling stimulates the OCR, a little more strongly in myoblasts than in myotubes, and inhibition of mitochondrial electron transfer diminishes it (Fig. 1A). Myocytes compensate for their inability to make ATP through oxidative phosphorylation by raising their rate of anaerobic glycolysis in response to oligomycin, as is clear from the increased proton production rate due to release of lactate^−^ and H^+^ (PPR_LAC_, Fig. 1B). Normalising OCR and PPR_LAC_ to number of myocyte nuclei allows statistical evaluation of differences in oxygen consumption and lactate-induced medium acidification between myoblasts and myotubes (Fig. 1C). Basal, ATP-synthesis-coupled and uncoupled respiration are all lower in myotubes than in myoblasts, but only the uncoupled rate difference is statistically significant. Neither respiration associated with proton leak nor *non*-mitochondrial oxygen consumption differs between differentiation states. Lactate-related medium acidification is somewhat higher in myotubes than myoblasts, particularly in the absence of respiratory effectors, but not significantly so (Fig. 1C). The proportion of myocyte respiration that is used to make ATP (Fig. 1D) is lower in myotubes (74%) than in myoblasts (84%). This difference in coupling efficiency of oxidative phosphorylation is statistically significant and is likely due to the higher workload faced by spontaneously contracting myotubes than by resting myoblasts, much akin to the comparably low fuel efficiency of car engines running at high speed. Indeed, low myotube coupling efficiency resembles the low value reported for intact perfused rat muscle [27]. Consistently, cellular respiratory control, i.e., the ratio of uncoupled and oligomycin-inhibited respiration, is lower in myotubes than myoblasts (Fig. 1E).

**Figure 1.**
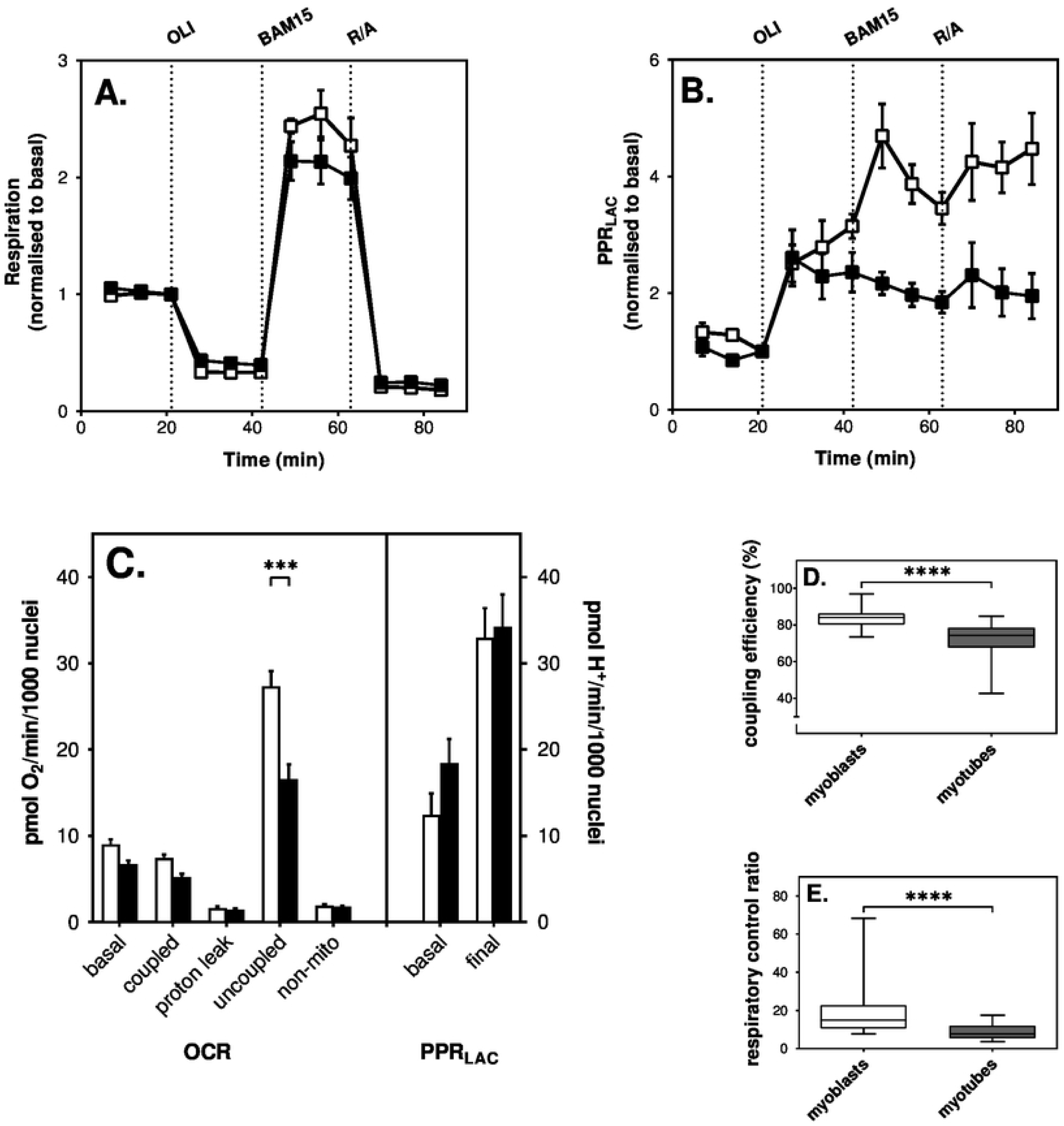
Myocellular respiration and proton release. Oxygen uptake and medium acidification by static L6 myoblasts (open symbols) and spontaneously contracting myotubes (filled symbols) were measured by extracellular flux (XF) analysis. Panels **A** and **B**: Respiratory and medium acidification traces, respectively, are based on the means ± SEM of 4 individual extracellular flux runs with the differentiation states measured 3-5 times in each. Medium acidification traces were corrected for CO_2_ contribution and thus reflect the proton production rate due to release of lactate– and H+ (PPR_LAC_) alone. Rates were normalised to the 3rd measurement and were obtained in the absence of any respiratory effector or in the cumulative presence of 5 µg/mL oligomycin (OLI), 1 µM uncoupler (BAM15), and a mixture of 1 µM rotenone and 1 µM antimycin A (R/A). Panel **C**: After normalisation to the number of myocyte nuclei, respiratory and proton production rates were used to calculate the basal oxygen consumption and medium acidification rates as well as the oxygen uptake activity that was coupled to ATP synthesis (coupled) or associated with mitochondrial proton leak. The maximum mitochondrial respiratory rate (uncoupled) and the rate of non-mitochondrial oxygen consumption (non-mito) are shown too, as is the PPR_LAC_ measured after addition of all effectors (final). Data are means ± SEM of 13-14 measurements sampled from 4 independent XF assays. Extracellular flux differences between myoblasts and myotubes were evaluated for statistical significance by 2-way ANOVA applying a Šídák’s multiple comparisons test (*** P < 0.001). Panels **D** and **E**: Coupling efficiencies and cell respiratory control ratios, respectively, were calculated from the data shown in Panel **C** and the control (0 µM NaNO_2_) data shown in Figure 4. Box-and-whiskers plots represent 25-34 measurements sampled from 8-9 independent XF assays. Statistical significance of the differences between myoblasts and myotubes (**** P < 0.0001) was evaluated by Mann Whitney tests.

From basal and oligomycin-sensitive OCR and basal PPR_LAC_ we then calculated absolute ATP supply rates, and found that total supply flux is marginally, but not statistically significantly, higher in myoblasts than in myotubes (Fig. 2). Mitochondrial ATP supply is also faster in myoblasts than myotubes, while glycolytic ATP supply tends to be slower (Fig. 2). When normalised to total ATP supply, the rate differences are statistically significant, which shows that L6 myocytes become more glycolytic after differentiation to contractile myotubes.

**Figure 2.**
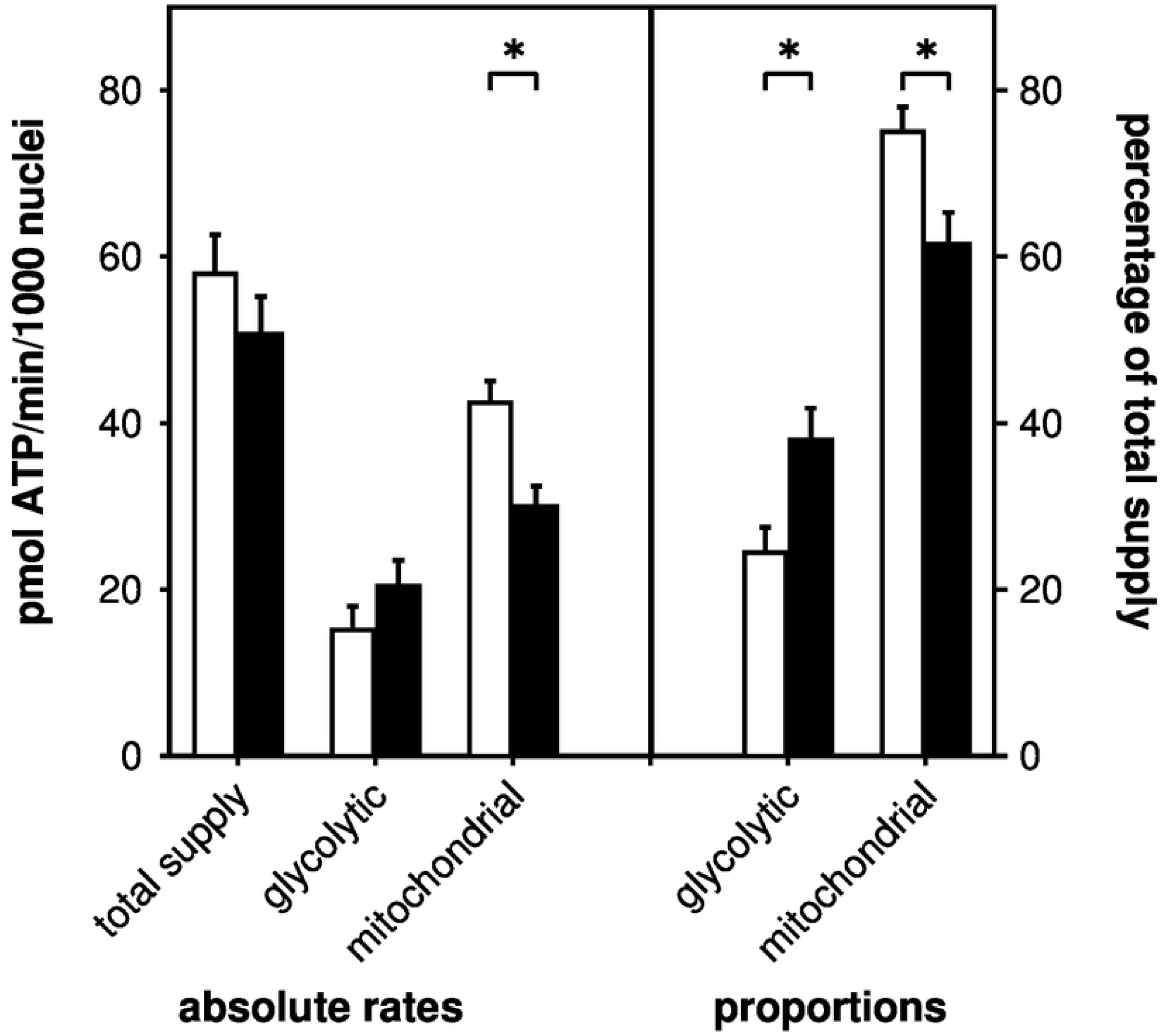
Myocellular ATP supply. Rates of glycolytic and mitochondrial ATP synthesis were calculated from data shown in Fig. 1C and were normalised to number of myocyte nuclei (absolute rates) or expressed as percentage of the combined, i.e., total ATP supply (proportions). Myoblast (open bars) and myotube (filled bars) data are means ± SEM of 13-14 measurements sampled from 4 separate XF assays. Bioenergetic differences between myoblasts and myotubes were evaluated for statistical significance by 2-way ANOVA applying a Šídák’s multiple comparisons test (* P < 0.05).

### Nitrite lowers coupling efficiency of oxidative phosphorylation

Seeking to establish how the bioenergetics of skeletal muscle cells are affected by nitrite, we exposed resting L6 myoblasts and spontaneously contracting myotubes to low-micromolar NaNO_2_ concentrations during the extracellular flux assays. Given the reported stimulation of oxidative ATP synthesis efficiency by dietary nitrate [15], we first measured possible NaNO_2_ effects on myocyte respiration. Linear regression analysis of such measurements shows that a ½-hour NaNO_2_ exposure is sufficient to raise the basal oxygen consumption of myoblasts in a dose-dependent manner (Fig. 3A). This NaNO_2_-induced respiratory rise is statistically significant and is due to stimulation of respiration linked to both ATP synthesis (Fig. 3B) and mitochondrial proton leak (Fig. 3C). NaNO_2_ does not affect uncoupled respiration (Fig. 3D) and lowers non-mitochondrial oxygen consumption (Fig. 3E). NaNO_2_ effects on respiration are less pronounced in myotubes than myoblasts, even when applied up to 5 instead of 1 µM (Figs 3F – 3J).

**Figure 3.**
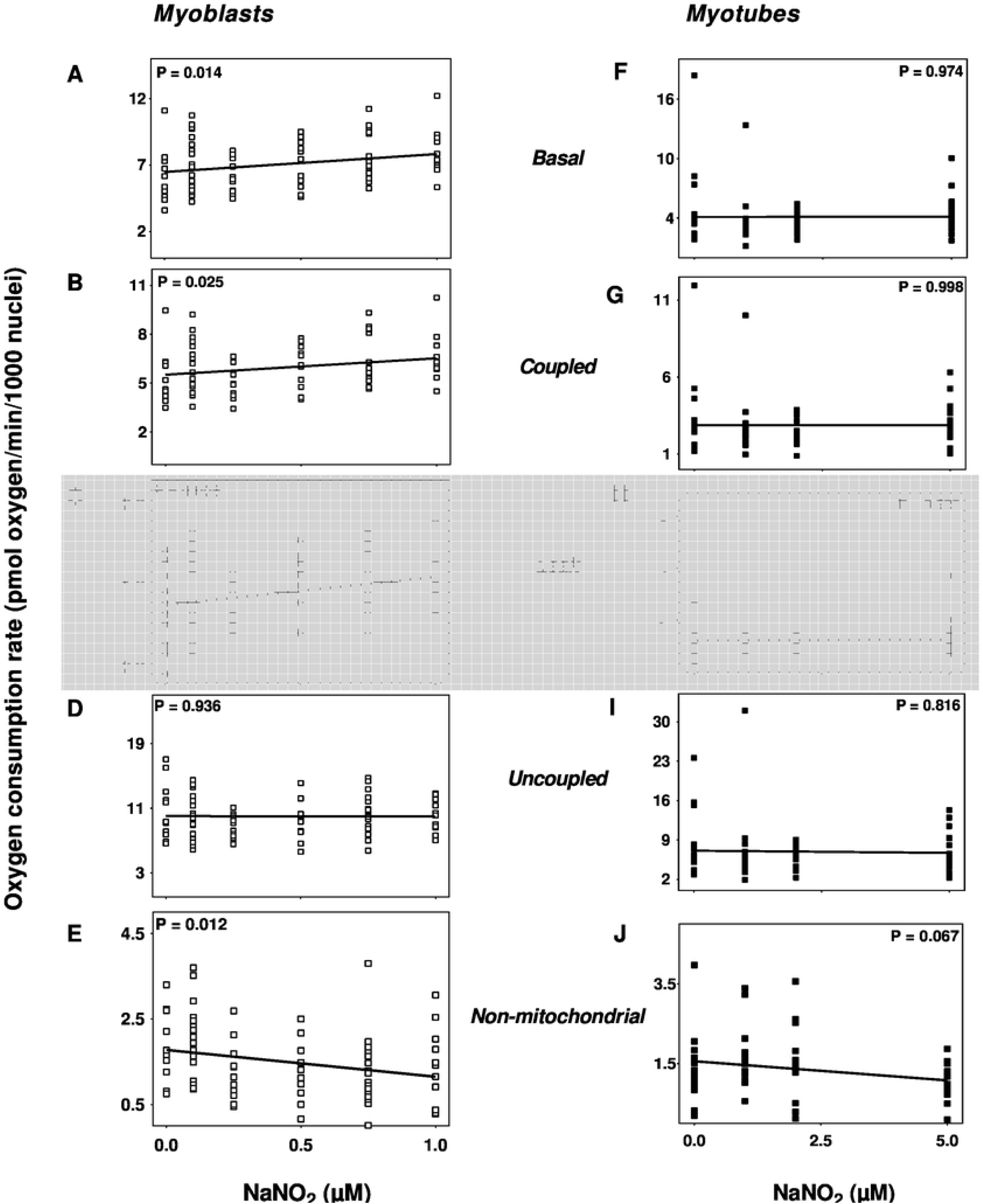
Nitrite effects on myocellular respiration. Mitochondrial and non-mitochondrial oxygen uptake rates were measured in static myoblasts (open symbols) and spontaneously contracting myotubes (filled symbols) after a ½-hour exposure to NaNO_2_ as described in Materials and Methods. Respiratory activities were normalised to number of myocyte nuclei and were obtained in the absence of effectors (Basal) or in the cumulative presence of 5 µg/mL oligomycin (Leak), 1 µM FCCP (Uncoupled) and a mix of 1 µM rotenone and 1 µM antimycin A (Non-mitochondrial). Oligomycin-sensitive oxygen uptake was used to estimate respiration coupled to ATP synthesis (Coupled). Data were obtained from 4-5 independent XF assays. NaNO_2_ effects were evaluated by linear regression analysis and a P value < 0.05 indicates the slope of the fitted data deviates significantly from zero.

As NaNO_2_ stimulates respiration linked to proton leak more strongly than respiration coupled to ATP synthesis, it attenuates coupling efficiency of oxidative phosphorylation, significantly so in myoblasts (Fig. 4A) and tending that direction in myotubes (Fig. 4B). Consistently, NaNO_2_ lowers the cell respiratory control ratios in a similar way (Figs 4C and 4D). Notably, NaNO_2_-induced attenuation of mitochondrial coupling efficiency is at odds with the reported stimulatory effect of NaNO_3_ on the P/O ratio of skeletal muscle mitochondria [15], but is consistent with the more recently reported lack of beneficial effect of dietary nitrate on mitochondrial efficiency [16,17].

**Figure 4.**
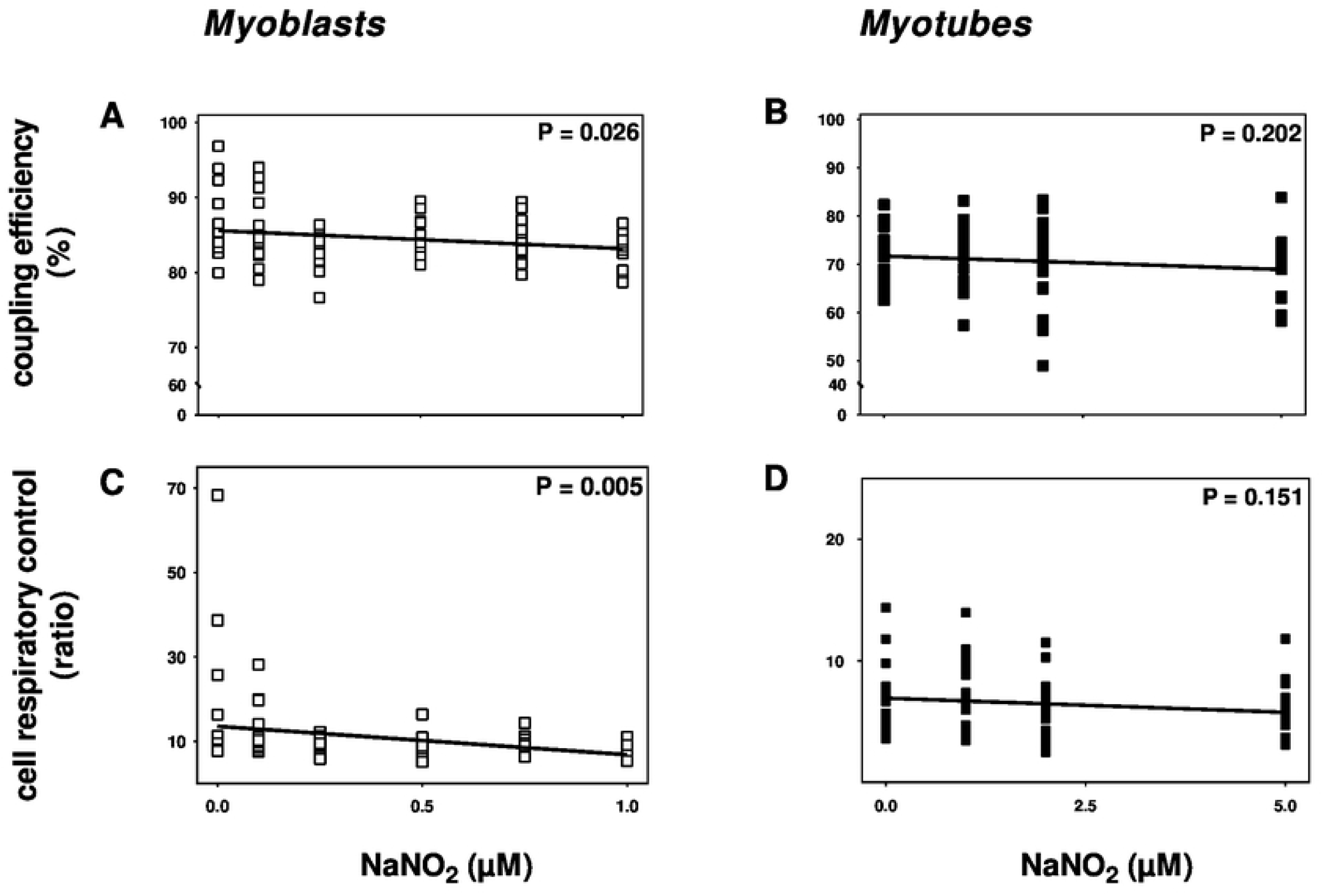
Nitrite lowers mitochondrial efficiency. Respiratory rates shown in Fig. 3 were used to calculate coupling efficiency of oxidative phosphorylation (Panels **A** and **B**) and cell respiratory control ratios (Panels **C** and **D**) in static myoblasts (open symbols) and spontaneously contracting myotubes (filled symbols). NaNO_2_ effects were evaluated by linear regression analysis and a P value < 0.05 indicates the slope of the fitted data deviates significantly from zero.

### Nitrite increases the rate of glycolytic and total ATP synthesis

Probing the attenuating effect of nitrite on mitochondrial coupling efficiency further, we next assessed whether NaNO_2_ also impeded the *rate* of oxidative phosphorylation. It transpires that NaNO_2_ does not significantly affect the rate of oxidative ATP synthesis in myotubes (Fig. 5B) and in fact increases it in myoblasts (Fig. 5A). Interestingly, calculating ATP supply flux from the XF data [21] revealed that NaNO_2_ raises the rate of glycolytic ATP supply both in myoblasts (Fig. 5C) and myotubes (Fig. 5D). Consistent with this surprising discovery, enzyme-based measurement of lactic acid release by myotubes during extracellular flux assays shows that NaNO_2_ increases the specific lactate production rate from 1.3 ± 0.52 to 3.1 ± 1.3 pmol/min/µg protein (P < 0.05). NaNO_2_ stimulation of glycolytic ATP supply flux is statistically significant in myoblasts and myotubes, either when rates are normalised to number of nuclei (Figs 5C and 5D) or to the overall ATP supply flux (Figs 5G and 5H). The total ATP supply is itself increased by NaNO_2_, an acute effect that is significant in myoblasts (Fig. 5E) and tends to significance in myotubes (Fig. 5F). Notably, NaNO_2_ has a highly statistically significant stimulatory effect on total ATP supply in both myoblasts (Fig. 5I) and myotubes (Fig. 5J) when this supply is normalised to basal cellular respiration. NaNO_2_ thus lowers the apparent oxygen cost of total ATP synthesis. Effects on myocellular ATP supply are specific to nitrite, as *nitrate* does not affect the rates of total, glycolytic or oxidative ATP supply in myoblasts or myotubes that were exposed for a ½ hour to NaNO_3_ (Fig. 6) at concentrations seen in circulation after dietary nitrate supplementation [5]. Notably, nitrate leaves the oxygen cost of ATP supply unaffected as well (Figs 6D and 6H).

**Figure 5.**
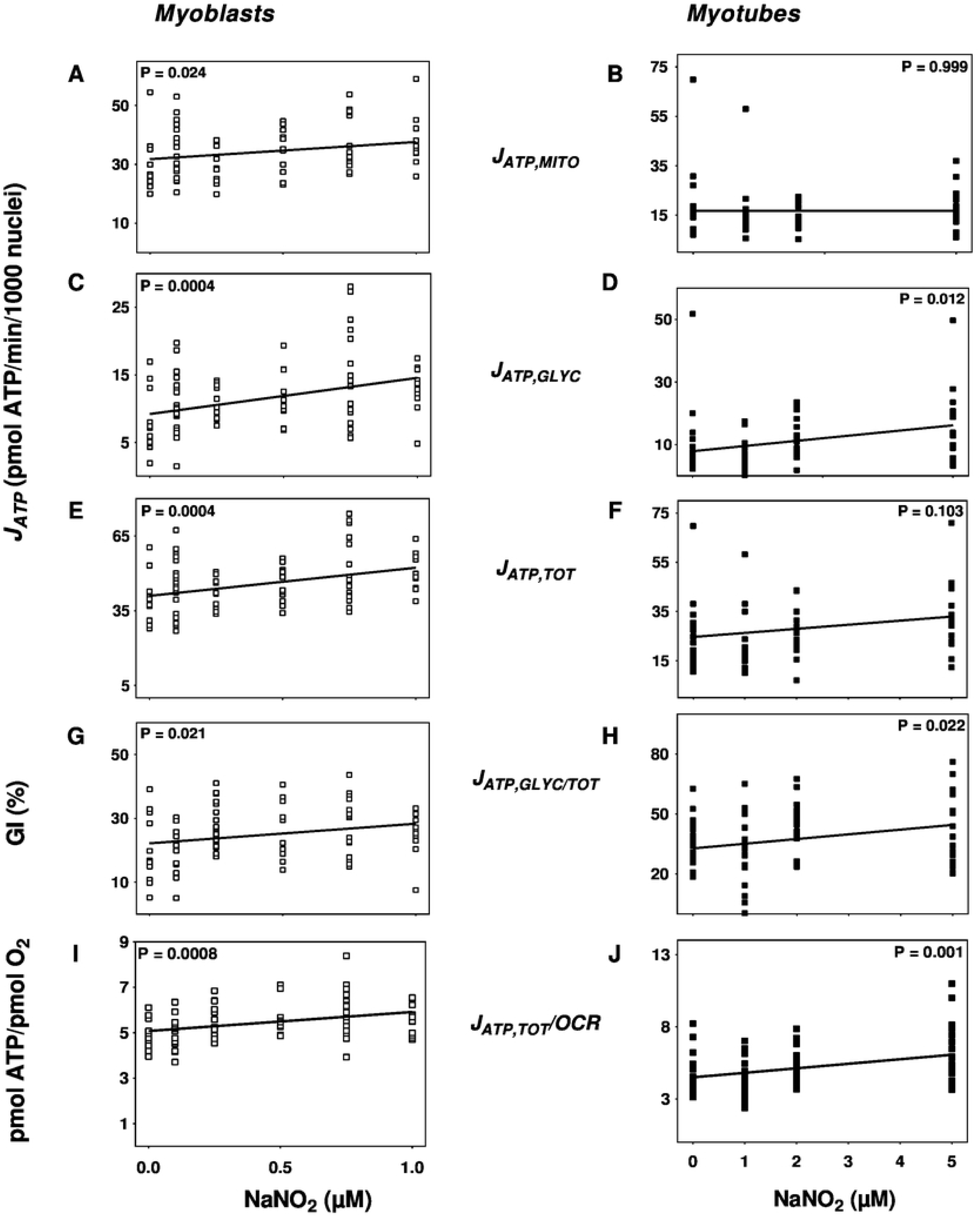
Nitrite effects on ATP supply. Respiratory rates shown in Fig. 3 were used, combined with concomitantly measured acidification rates (*cf*. Figs 1B, 1C), to calculate rates of mitochondrial (Panels **A** and **B**), glycolytic (Panels **C** and **D**) and total (Panels **E** and **F**) ATP supply (J_ATP,MITO_, J_ATP,GLYC_ and J_ATP,TOT_, respectively) in static myoblasts (open symbols) and spontaneously contracting myotubes (filled symbols). Glycolytic ATP synthesis rates are also shown as percentages of total ATP supply rate (J_ATP,GLY/TOT_, Panels **G** and **H**), i.e., as myocellular glycolytic indices (GI). Furthermore, total ATP synthesis rates are normalised to the total cellular oxygen consumption rate (J_ATP,TOT_/OCR, Panels **I** and **J**). NaNO_2_ effects were evaluated by linear regression analysis and a P value < 0.05 indicates the slope of the fitted data deviates significantly from zero.

**Figure 6.**
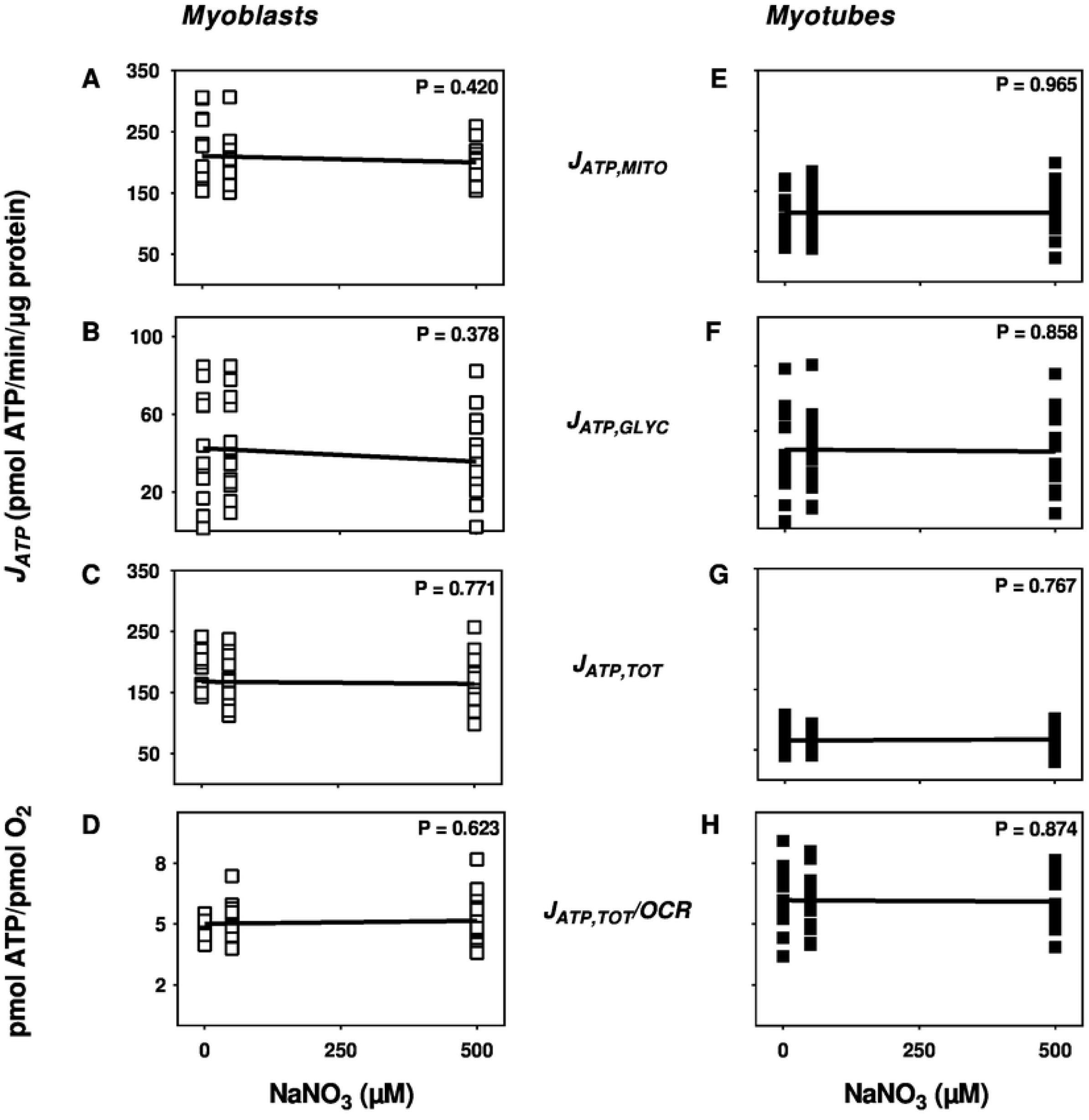
Nitrate effects on ATP supply. Rates of mitochondrial, glycolytic and total ATP supply (J_ATP,MITO_, J_ATP,GLYC_ and J_ATP,TOT_, respectively) in static myoblasts (open symbols) and spontaneously contracting myotubes (filled symbols) were calculated from oxygen uptake and medium acidification data obtained after a ½-hour exposure to NaNO_3_. Data were obtained from 4 separate XF assays. NaNO_3_ effects were evaluated by linear regression analysis. High P values indicate fitted data slopes do not deviate significantly from zero.

### Nitrite increases the cellular glycolytic index

The NaNO_2_ effect on myocellular ATP supply can be illustrated by ‘bioenergetic space’ plots that directly relate mitochondrial and glycolytic ATP synthesis rates [21]. Cells that make most of their ATP via oxidative metabolism occupy space above the main diagonal (from the origin with a slope of 1) of such plots (Fig. 7A, clear area), and have a glycolytic index (GI) [21] that is lower than 50%. Cells that rely predominantly on glycolysis for their ATP supply occupy space below the diagonal (Fig. 7A, shaded area) and have a GI higher than 50%. GI is defined as the percentage of total ATP supply from glycolysis (*cf*. Figs 5G and 5H) and is inversely related to the slope of diagonals that link the origin of the space plot to the position that reflects particular metabolic behaviour (Figs 7B and 7C, dotted lines).

**Figure 7.**
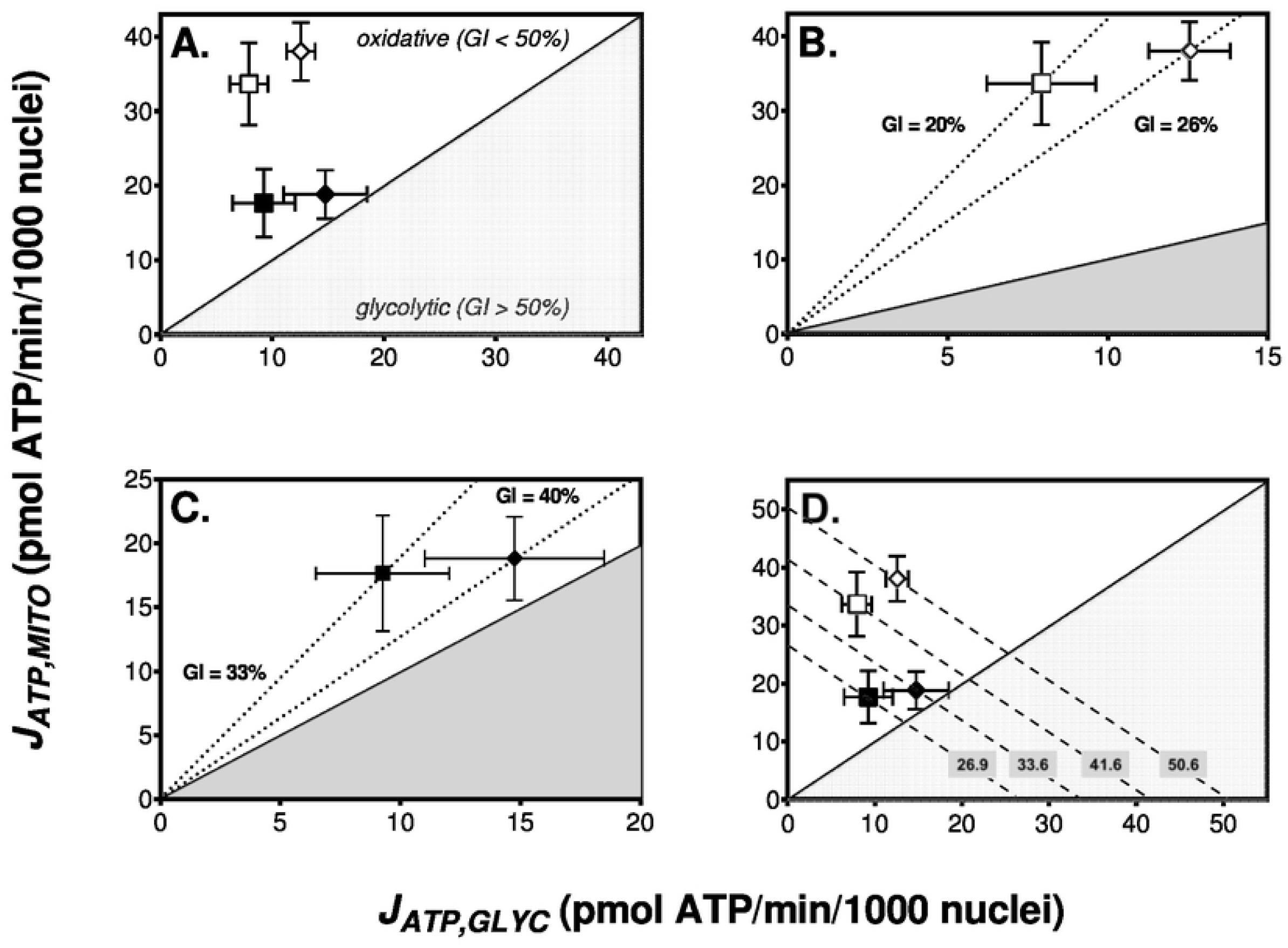
Nitrite increases the dependence of skeletal muscle cells on glycolytic ATP supply. Bioenergetic space plots [21] relate the rate of mitochondrial ATP supply (*J*_*ATP,MITO*_) to the rate of glycolytic ATP supply (*J*_*ATP,GLYC*_). Individual data obtained from experiments with static myoblasts (open symbols) and spontaneously contracting myotubes (filled symbols), in the absence (squares) or the presence (diamonds) of NaNO_2_ (1 and 5 µM for myoblasts and myotubes, respectively), were sourced from Fig. 5. Diagonal lines with positive slopes reflect the glycolytic index (GI), i.e., the percentage of total ATP synthesis from glycolysis. The white area above the GI_50%_ diagonal covers ‘oxidative’ bioenergetic space, whereas the grey shaded area below this diagonal covers the ‘glycolytic’ space. GI values indicated by the dotted lines in Panels **B** and **C** were calculated from the shown mean rates of oxidative and glycolytic ATP supply. The dashed diagonal lines with a slope of – 1 in Panel **D** indicate space with the same total rate of ATP supply. The numbers labelling these lines are mean supply rates in pmol ATP/min/1000 nuclei.

Myoblasts and myotubes assayed with 5 mM glucose as sole metabolic fuel both have a GI below 50% (Fig. 7A), and can thus be considered oxidative systems with most of their ATP made by mitochondria. Consistent with increased myocyte reliance on glycolytic ATP supply after differentiation (*cf*. Fig. 2), the mean GI of myoblasts (Fig. 7B – 20%) is lower than that of myotubes (Fig. 7C – 33%). NaNO_2_ pushes the bioenergetic behaviour of both systems acutely towards a more glycolytic phenotype, raising the GI of myoblasts to 26% on average (Fig. 7B) and that of myotubes to 40% (Fig. 7C). These GI effects are statistically significant when NaNO_2_ dose dependencies are taken into account (*cf*. Figs 5G and 5H, respectively).

Diagonals in bioenergetic space plots with slopes of –1 link positions at which the rate of total ATP supply is identical, but is accounted for by differential relative contributions of glycolytic and oxidative ATP supply flux (Fig. 7D, dashed lines). For instance, the total ATP supply is faster in myoblasts than in myotubes – even though the contractile differentiated L6 myocytes are more glycolytic (Fig. 7C) than their non-differentiated static counterparts (Fig. 7B) – and the respective steady states are thus described by separate negative diagonals. NaNO_2_-induced GI increments shift both steady states onto different diagonals (Fig. 7D), which confirms NaNO_2_ has increased total ATP supply, and glycolytic ATP synthesis has not been raised by NaNO_2_ at the expense of oxidative phosphorylation. NaNO_2_ lifts the rate of total ATP synthesis from 41.6 to 50.6 pmol ATP/min/1000 nuclei in myoblasts, and from 26.9 to 33.6 pmol ATP/min/1000 nuclei in myotubes. Taking dose dependencies into account, this NaNO_2_ effect is statistically significant in myoblasts (*cf*. Fig. 5E) and tends to significance in myotubes (*cf*. Fig. 5F). Consistent with the data shown in Fig. 2, total ATP supply rate tend to be lower in myotubes than myoblasts, irrespective of variation between experiments.

### Nitrite does not affect glucose uptake

When the bioenergetic behaviour of L6 myocytes is measured in buffer lacking metabolic fuel, injection of 2 mM glucose instantly stimulates acidification of the extracellular medium, but leaves oxygen consumption unaffected [22]. Such responses suggest the possibility that the NaNO_2_-induced increase in the glycolytic ATP synthesis rate is secondary to increased glucose availability. Indeed, nitrite has previously been reported to increase glucose uptake by cultured adipocytes [28], and we have seen acute NaNO_2_ stimulation of insulin-sensitive glucose uptake by L6 myocytes (Rosie Donnell and Charles Affourtit, unpublished data). Because the regulation of cellular glucose uptake, for example by insulin, depends on the cells’ exact growth history and assay conditions [29], we measured accumulation of 2-deoxyglucose in myocytes cultured in the same way as those used for the bioenergetic studies, and applied assay conditions that were near-identical. Glucose uptake is unaffected by a ½-hour NaNO_2_ exposure in myocytes treated in this representative manner (Fig. 8). The lack of glucose uptake phenotype holds for resting, insulin-unresponsive myoblasts as well as for spontaneously contracting, insulin-sensitive myotubes.

**Figure 8.**
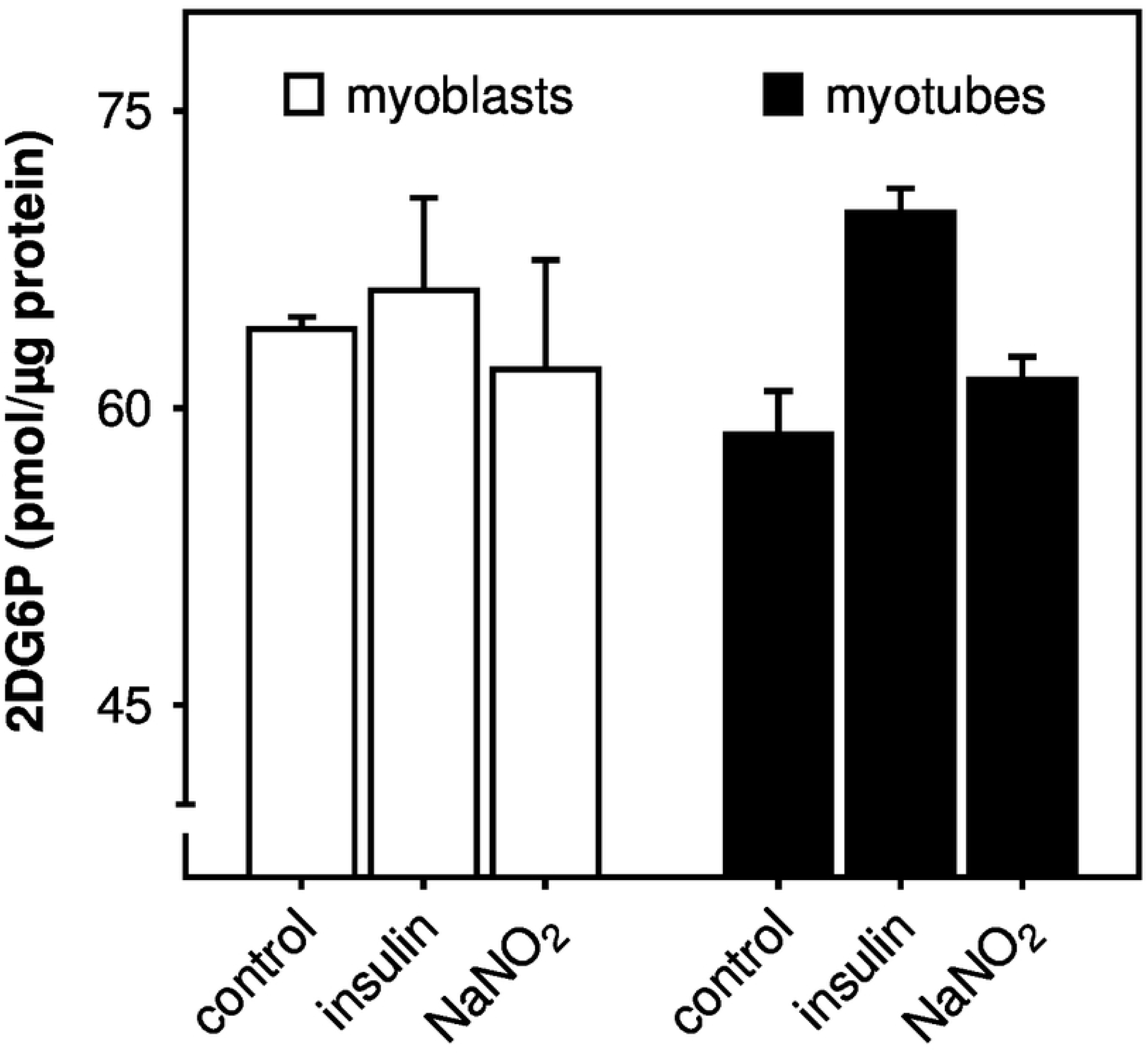
Nitrite does not affect glucose uptake by skeletal muscle cells. Static myoblasts (open bars) and spontaneously contracting myotubes (filled bars) were grown and assayed under the same conditions as those applied during extracellular flux analysis, and were exposed to 1 µM NaNO_2_ for 30 min with and without 100 nM human insulin. Glucose uptake was assayed as 2-deoxyglucose-6-phosphate (2DG6P) accumulated over a 30-min period. Data are means ± SEM of 3 independent experiments with each condition assayed in triplicate. Differences between differentiation state and assay conditions were evaluated by 2-way ANOVA and were found not significant.

## Discussion

The extracellular flux data reported in this paper highlight how simultaneous measurement of oxygen consumption and medium acidification by cultured cells can offer valuable insight in cellular energy metabolism when such data are used to quantify ATP supply flux. We thus reveal that the rate of glycolytic ATP synthesis in skeletal muscle cells is increased by a ½-hour NaNO_2_ exposure. This acute stimulation of glycolytic ATP supply does not occur at the expense of mitochondrial ATP synthesis, which means NaNO_2_ raises the overall rate of ATP supply, and pushes the myocytes towards a more glycolytic bioenergetic phenotype. As the increase in the overall ATP synthesis rate emerges against a relatively stable oxygen uptake background, NaNO_2_ lowers the apparent oxygen cost of ATP supply. This effect is not clear from the ‘raw’ oxygen uptake and medium acidification data. It remains uncertain at present how well our rat myocyte data will translate to human muscle physiology as the conditions of our *in vitro* experiments are clearly not identical to those found *in vivo*. The applied ambient oxygen tension will for example be much higher than the one that prevails in resting and contracting skeletal muscle. Moreover, the physiological relevance of *rodent* myocytes is of course limited, but it is worth emphasis that L6 cells have proven a reliable model of skeletal muscle bioenergetics in our hands before, with behaviour that was highly reproducible in primary human myocytes [22,30]. With these caveats, our findings inform ongoing debate as to the mechanism by which dietary nitrate alters muscle function.

### Oxygen cost of exercise

Dietary nitrate has been reported to lower the oxygen cost of human exercise by decreasing proton leak in skeletal muscle mitochondria, thus increasing coupling efficiency of oxidative phosphorylation [15]. Lowered proton leak was attributed to decreased protein expression of mitochondrial carriers whose contribution to proton leak is contentious, particularly under the bioenergetic conditions that prevail during exercise [14]. More recent human work suggests that lowered whole-body oxygen uptake may occur without mitochondrial efficiency changes [16,17], a notion supported by research on various model systems [31-34]. Our data add to this body of corroborating support by showing that nitrite – whose circulating [5] and intra-myocellular [20] concentrations are raised after dietary nitrate exposure – does *not* increase coupling efficiency of oxidative phosphorylation in myoblasts (Fig. 4A) or in spontaneously contracting myotubes (Fig. 4B). In fact, nitrite acutely lowers coupling efficiency in myoblasts and tends to do so in myotubes.

Human and rodent studies have shown that nitrate increases contractile efficiency of skeletal muscle [35,36]. Such a nitrate effect lowers the ATP cost of muscle work and may thus be expected to benefit high intensity work the same as, if not more than, low intensity work. The relation between nitrate benefit and muscle workload, however, appears the opposite: nitrate lowers the oxygen cost of sub-maximal intensity exercise [37] and may even lower resting metabolic rate [38], but leaves severe or maximal intensity exercise unaffected [12]. Our data suggest an alternative explanation for the nitrate exercise phenotype, i.e., a rise in the myocellular glycolytic index (Fig. 7). The raised GI follows from a nitrite-induced stimulation of glycolytic ATP supply, which occurs without loss of oxidative ATP supply. Mitochondrial oxygen uptake changes relatively little following nitrite exposure, whilst non-mitochondrial respiration decreases (Fig. 3), which thus lowers the oxygen cost of ATP synthesis (Fig. 5). Notably, this apparent oxygen cost benefit does not reflect increased efficiency of oxidative metabolism, which is actually attenuated (Fig. 4). Our results suggest the possibility, albeit speculative at present, that dietary nitrate fails to lower the oxygen cost of severe and maximal intensity exercise, because augmentation of an already high lactate output [39,40] accrues additional oxygen demand that cancels out any respiratory benefit. Such oxygen demand reflects the energetic cost of lactate recycling through hepatic gluconeogenesis [41], which is repaid in recovery and perhaps also during exercise. Our findings appear at odds with the reported lack of nitrate effect on circulating lactate [8]. Effects of nitrate on plasma lactate concentration are variable though, and improved muscle function may also coincide with increased [39,40,42] and decreased [43] lactate levels. More work is required to clarify this variation. In this respect, the circulating lactate concentration is perhaps not the most suitable metric to quantify skeletal muscle lactate release *in vivo*, as this concentration is set by other processes too, for example, hepatic gluconeogenesis [41].

Dietary nitrate effects on skeletal muscle function depend on fibre type composition [44]. For instance, nitrate increases oxygen perfusion [45] and calcium handling [36] of murine fast-twitch (type II) but not slow-switch (type I) fibres. Consistently, nitrate-lowering effects on the systemic oxygen uptake rate are most pronounced during exercise where type II fibres are recruited predominantly [46-48]. Nitrate selectivity is possibly related to the relatively low microvascular oxygen tension of type II fibres [49], as exercise benefits are more prominent in hypoxia [50,51] than normoxia [52]. In this respect, it is worth emphasis that type II fibres are glycolytic systems [53] that perhaps benefit more than oxidative type I fibres [53] from the nitrite-induced stimulation of glycolytic ATP supply reported here (Fig. 7).

### Mechanism of bioenergetic nitrite effect

Our results do not reveal the mechanism by which nitrite stimulates glycolytic ATP supply, but they exclude several possibilities. Changes in mitochondrial and glycolytic ATP synthesis are often reciprocal. For instance, an increased glycolytic ATP synthesis rate may arise as compensatory response to impeded mitochondrial ATP supply (Pasteur effect – *cf*. Figs 1A and 1B for opposite effects of oligomycin on respiration and proton production). Instead, an increase in glycolytic ATP supply may suppress oxidative ATP synthesis (Crabtree effect). However, it is unlikely that our nitrite phenotype reflects such ‘bioenergetic supply flexibility’ [21], as stimulation of glycolytic ATP supply does not coincide with lowered mitochondrial respiration. Although nitrite lowers efficiency of oxidative phosphorylation (Fig. 4), it does not lower the rate (Fig. 5). The maintained rate of oxidative phosphorylation renders it unlikely that cytochrome *c* oxidase has been inhibited by nitrite-derived nitric oxide [54]. Dietary nitrate effects on glucose homeostasis [55] may be related to increased skeletal muscle glucose disposal [56]. Increased myocellular glucose availability could, in principle, explain the bioenergetic nitrite effect we report here (*cf*. [22]), but such an explanation is weakened by the lack of nitrite effect on 2-deoxyglucose uptake (Fig. 8). Increased glucose availability could also be achieved by nitrite stimulation of glycogenolysis, a possibility that cannot be excluded by our data.

It is conceivable that glycolytic ATP supply is raised indirectly in response to increased ATP turnover as skeletal muscle ATP flux is predominantly demand-driven [57]. However, it is unlikely that nitrite stimulation of energy expenditure is due to increased contractile activity, as we observe the glycolytic nitrite phenotype both in spontaneously contracting myotubes and resting myoblasts. If nitrite indeed stimulates ATP consumption, then it will also have to be explained why raised ATP demand is mostly met by glycolytic rather than mitochondrial ATP supply. In addition to, or instead of, indirect stimulation of glycolysis, it is of course also possible that the activity of one or more glycolytic enzymes is modulated directly by nitrite. Although the nitrite phenotype emerges on a relatively short timescale, such modulation may involve upregulation of gene expression.

Irrespective of the site(s) of action, it also remains to be established whether the nitrite effect is direct or mediated by a nitrite-derivative. Whilst nitric oxide is held responsible for dietary nitrate protection against various pathologies [58], it remains uncertain if this radical species is involved in the human exercise phenotype [14,59]. The low pH and low oxygen tensions required to reduce nitrite are unlikely to be present in the skeletal muscle cells of healthy subjects [14]. Indeed, the acute nitrite effect on the oxygen cost of ATP synthesis reported here is achieved at neutral pH and under an oxygen atmosphere (21%) that is hyperoxic from a physiological perspective. Under these conditions, it is unlikely that nitrite is reduced to nitric oxide.

### Concluding remarks

The quantitative analysis of myocellular bioenergetics reported here sheds fresh light on the elusive mechanism that underlies the exercise benefit of nitrate. The impact of our findings on human physiology remains to be confirmed, but we feel that the reported data contribute towards the mechanistic understanding that will be crucial for achieving the full translational potential of dietary nitrate supplementation.

## Acknowledgements

The reported work was supported by a BBSRC-funded Daphne Jackson Trust Fellowship to AGW with consumable support from the University of Plymouth. The Daphne Jackson Trust, BBSRC nor the University of Plymouth were involved in the design and execution of this study, the analysis and interpretation of the data or in the writing of the manuscript. It was the decision of the authors only to submit the manuscript for publication. We thank Drs Rosie Donnell, Jane Carré (University of Plymouth), Andy Jones and Paul Winyard (University of Exeter) for stimulating discussion.

## References

1. Mills CE, Khatri J, Maskell P, Odongerel C, Webb AJ. It is rocket science - why dietary nitrate is hard to Beet! part II: further mechanisms and therapeutic potential of the nitrate-nitrite-NO pathway. Br J Clin Pharmacol 2017;83: 140–151.

2. Jackson JK, Patterson AJ, MacDonald-Wicks LK, Oldmeadow C, McEvoy MA. The role of inorganic nitrate and nitrite in cardiovascular disease risk factors: a systematic review and meta-analysis of human evidence. Nutrition Reviews 2018;76: 348–371.

3. Lundberg JO, Carlström M, Weitzberg E. Metabolic effects of dietary nitrate in health and disease. Cell Metab 2018;28: 9–22.

4. Duncan C, Dougall H, Johnston P, Green S, Brogan R, Leifert C, et al. Chemical generation of nitric oxide in the mouth from the enterosalivary circulation of dietary nitrate. Nat Med 1995;1: 546–551.

5. Webb AJ, Patel N, Loukogeorgakis S, Okorie M, Aboud Z, Misra S, et al. Acute blood pressure lowering, vasoprotective, and antiplatelet properties of dietary nitrate via bioconversion to nitrite. Hypertension 2008;51: 784–790.

6. Li H, Samouilov A, Liu X, Zweier JL. Characterization of the magnitude and kinetics of xanthine oxidase-catalyzed nitrate reduction: evaluation of its role in nitrite and nitric oxide generation in anoxic tissues. Biochemistry 2003;42: 1150–1159.

7. Cosby K, Partovi KS, Crawford JH, Patel RP, Reiter CD, Martyr S, et al. Nitrite reduction to nitric oxide by deoxyhemoglobin vasodilates the human circulation. Nat Med 2003;9: 1498–1505.

8. Larsen FJ, Weitzberg E, Lundberg JO, Ekblom B. Effects of dietary nitrate on oxygen cost during exercise. Acta Physiol 2007;191: 59–66.

9. Bailey SJ, Winyard P, Vanhatalo A, Blackwell JR, Dimenna FJ, Wilkerson DP, et al. Dietary nitrate supplementation reduces the O_2_ cost of low-intensity exercise and enhances tolerance to high-intensity exercise in humans. J Appl Physiol 2009;107: 1144–1155.

10. Poole DC, Richardson RS. Determinants of oxygen uptake. Implications for exercise testing. Sports Med 1997;24: 308–320

11. Jones AM, Thompson C, Wylie LJ, Vanhatalo A. Dietary nitrate and physical performance. Annu Rev Nutr 2018;38: 303–328.

12. Pawlak-Chaouch M, Boissière J, Gamelin FX. Effect of dietary nitrate supplementation on metabolic rate during rest and exercise in human: A systematic review and a meta-analysis. Nitric Oxide 2016;53: 65–76.

13. McMahon NF, Leveritt MD, Pavey TG. The effect of dietary nitrate supplementation on endurance exercise performance in healthy adults: a systematic review and meta-analysis. Sports Med 2017;47: 735–756.

14. Affourtit C, Bailey SJ, Jones AM, Smallwood MJ, Winyard PG. On the mechanism by which dietary nitrate improves human skeletal muscle function. Front Physiol 2015;6: 211.

15. Larsen FJ, Schiffer TA, Borniquel S, Sahlin K, Ekblom B, Lundberg JO, et al. Dietary inorganic nitrate improves mitochondrial efficiency in humans. Cell Metab. 2011;13: 149–159.

16. Whitfield J, Ludzki A, Heigenhauser GJF, Senden JMG, Verdijk LB, van Loon LJC, et al. Beetroot juice supplementation reduces whole body oxygen consumption but does not improve indices of mitochondrial efficiency in human skeletal muscle. J Physiol 2016;594: 421–435.

17. Ntessalen M, Procter NEK, Schwarz K, Loudon BL, Minnion M, Fernandez BO, et al. Inorganic nitrate and nitrite supplementation fails to improve skeletal muscle mitochondrial efficiency in mice and humans. Am J Clin Nutr 2020;111: 79–89.

18. Coggan AR, Peterson LR. Dietary nitrate enhances the contractile properties of human skeletal muscle. Exerc Sport Sci Rev 2018;46: 254–261.

19. Kemp GJ. Beetroot juice supplementation reduces the oxygen cost of exercise without improving mitochondrial efficiency: but how? J Physiol 2016;594: 253.

20. Srihirun S, Park JW, Teng R, Sawaengdee W, Piknova B, Schechter AN. Nitrate uptake and metabolism in human skeletal muscle cell cultures. Nitric Oxide 2020;94: 1–8.

21. Mookerjee SA, Gerencser AA, Nicholls DG, Brand MD. Quantifying intracellular rates of glycolytic and oxidative ATP production and consumption using extracellular flux measurements. J Biol Chem 2017;292: 7189–7207.

22. Nisr RB, Affourtit C. Insulin acutely improves mitochondrial function of rat and human skeletal muscle by increasing coupling efficiency of oxidative phosphorylation. Biochim Biophys Acta. 2014;1837: 270–276.

23. Gerencser AA, Neilson A, Choi SW, Edman U, Yadava N, Oh RJ, et al. Quantitative microplate-based respirometry with correction for oxygen diffusion. Anal Chem 2009;81: 6868–6878.

24. Barlow J, Hirschberg-Jensen V, Affourtit C. Uncoupling protein-2 attenuates palmitoleate protection against the cytotoxic production of mitochondrial reactive oxygen species in INS-1E insulinoma cells. Redox Biol 2015;4: 14–22.

25. Brand MD, Nicholls DG. Assessing mitochondrial dysfunction in cells. Biochem J 2011;435: 297–312.

26. Mookerjee SA, Goncalves RLS, Gerencser AA, Nicholls DG, Brand MD. The contributions of respiration and glycolysis to extracellular acid production. Biochim Biohys Acta 2015;1847: 171–181.

27. Rolfe DF, Brand MD. Contribution of mitochondrial proton leak to skeletal muscle respiration and to standard metabolic rate. Am J Physiol 1996;271: C1380–1389.

28. Khoo NKH, Mo L, Zharikov S, Kamga-Pride C, Quesnelle K, Golin-Bisello F, et al. Nitrite augments glucose uptake in adipocytes through the protein kinase A-dependent stimulation of mitochondrial fusion. Free Radic Biol Med 2014;70: 45–53.

29. Ching JK, Rajguru P, Marupudi N, Banerjee S, Fisher JS. A role for AMPK in increased insulin action after serum starvation. Am J Physiol 2010;299: C1171–1179.

30. Nisr RB, Affourtit C. Palmitate-induced changes in energy demand cause reallocation of ATP supply in rat and human skeletal muscle cells. Biochim Biophys Acta 2016;1857: 1403–1411.

31. Ivarsson N, Schiffer TA, Hernández A, Lanner JT, Weitzberg E, Lundberg JO, et al. Dietary nitrate markedly improves voluntary running in mice. Physiol Behav 2017;168: 55–61.

32. Monaco CMF, Miotto PM, Huber JS, van Loon LJC, Simpson JA, Holloway GP. Sodium nitrate supplementation alters mitochondrial H2O2 emission but does not improve mitochondrial oxidative metabolism in the heart of healthy rats. Am J Physiol 2018;315: R191–R204.

33. Axton ER, Beaver LM, St Mary L, Truong L, Logan CR, Spagnoli S, et al. Treatment with nitrate, but not nitrite, lowers the oxygen cost of exercise and decreases glycolytic intermediates while increasing fatty acid metabolites in exercised zebrafish. J Nutr 2019;149: 2120–2132.

34. Porcelli S, Rasica L, Ferguson BS, Kavazis AN, McDonald J, Hogan MC, et al. Effect of acute nitrite infusion on contractile economy and metabolism in isolated skeletal muscle in situ during hypoxia. J Physiol 2020;598: 2371–2384.

35. Bailey SJ, Fulford J, Vanhatalo A, Winyard PG, Blackwell JR, DiMenna FJ, et al. Dietary nitrate supplementation enhances muscle contractile efficiency during knee-extensor exercise in humans. J Appl Physiol 2010;109: 135–148.

36. Hernández A, Schiffer TA, Ivarsson N, Cheng AJ, Bruton JD, Lundberg JO, et al. Dietary nitrate increases tetanic [^2+^]_i_ and contractile force in mouse fast-twitch muscle. J Physiol 2012;590: 3575–3583.

37. Pawlak-Chaouch M, Boissière J, Gamelin FX, Cuvelier G, Berthoin S, Aucouturier Effect of dietary nitrate supplementation on metabolic rate during rest and exercise in human: A systematic review and a meta-analysis. Nitric Oxide. 2016;53: 65–76.

38. Larsen FJ, Schiffer TA, Ekblom B, Mattsson MP, Checa A, Wheelock CE, et al. Dietary nitrate reduces resting metabolic rate: a randomized, crossover study in humans. Am J CliN Nutr 2014;99: 843–850.

39. Domínguez R, Garnacho-Castaño MV, Cuenca E, García-Fernández P, Muñoz-González A, de Jesús F, et al. Effects of beetroot juice supplementation on a 30-s high-intensity inertial cycle ergometer test. Nutrients 2017;9: 1360.

40. Shannon OM, Barlow MJ, Duckworth L, Williams E, Wort G, Woods D, et al. Dietary nitrate supplementation enhances short but not longer duration running time-trial performance. Eur J Appl Physiol. 2017;117: 775–785.

41. Waterhouse C, Keilson J. Cori cycle activity in man. J Clin Invest 1969;48: 2359–2366.

42. Wylie LJ, Bailey SJ, Kelly J, Blackwell JR, Vanhatalo A, Jones AM. Influence of beetroot juice supplementation on intermittent exercise performance. Eur J Appl Physiol 2016;116: 415–425.

43. Santana J, Madureira D, de França E, Rossi F, Rodrigues B, Fukushima A, et al. Nitrate supplementation combined with a running training program improved time-trial performance in recreationally trained runners. Sports 2019;7: 120.

44. Jones AM, Ferguson SK, Bailey SJ, Vanhatalo A, Poole DC. Fiber Type-Specific Effects of Dietary Nitrate. Exerc Sport Sci Rev 2016;44: 53–60.

45. Ferguson SK, Holdsworth CT, Wright JL, Fees AJ, Allen JD, Jones AM, et al. Microvascular oxygen pressures in muscles comprised of different fiber types: Impact of dietary nitrate supplementation. Nitric Oxide 2014;48: 38–43.

46. Breese BC, McNarry MA, Marwood S, Blackwell JR, Bailey SJ, Jones AM. Beetroot juice supplementation speeds O_2_ uptake kinetics and improves exercise tolerance during severe-intensity exercise initiated from an elevated metabolic rate. Am J Physiol 2013;305: R1441–R1450.

47. Bailey SJ, Varnham RL, Dimenna FJ, Breese BC, Wylie LJ, Jones AM. Inorganic nitrate supplementation improves muscle oxygenation, O_2_ uptake kinetics, and exercise tolerance at high but not low pedal rates. J Appl Physiol 2015;118: 1396–1405.

48. Coggan AR, Leibowitz JL, Kadkhodayan A, Thomas DP, Ramamurthy S, Spearie CA, et al. Effect of acute dietary nitrate intake on maximal knee extensor speed and power in healthy men and women. Nitric Oxide 2015;48: 16–21.

49. McDonough P, Behnke BJ, Padilla DJ, Musch TI, Poole DC. Control of microvascular oxygen pressures in rat muscles comprised of different fibre types. J Physiol 2005;563: 903–913.

50. Masschelein E, Van Thienen R, Wang X, Van Schepdael A, Thomis M, Hespel P. Dietary nitrate improves muscle but not cerebral oxygenation status during exercise in hypoxia. J Appl Physiol 2012;113: 736–745.

51. Muggeridge DJ, Howe CCF, Spendiff O, Pedlar C, James PE, Easton C. A single dose of beetroot juice enhances cycling performance in simulated altitude. Med Sci Sports Exerc 2014;46: 143–150.

52. Kelly J, Vanhatalo A, Bailey SJ, Wylie LJ, Tucker C, List S, et al. Dietary nitrate supplementation: effects on plasma nitrite and pulmonary O2 uptake dynamics during exercise in hypoxia and normoxia. Am J Physiol 2014;307: R920–R930.

53. Bottinelli R, Reggiani C. Human skeletal muscle fibres: molecular and functional diversity. Progr Biophys Mol Biol 2000;73: 195–262.

54. Brown G, Borutaite V. Nitric oxide and mitochondrial respiration in the heart. Cardiovasc Res 2007;75: 283–290.

55. Carlström M, Larsen FJ, Nyström T, Hezel M, Borniquel S, Weitzberg E, et al. Dietary inorganic nitrate reverses features of metabolic syndrome in endothelial nitric oxide synthase-deficient mice. Proc Natl Acad Sci 2010;107: 17716–17720.

56. Jiang H, Torregrossa AC, Potts A, Pierini D, Aranke M, Garg HK, et al. Dietary nitrite improves insulin signaling through GLUT4 translocation. Free Radic Biol Med 2014;67: 51–57.

57. Affourtit C. Mitochondrial involvement in skeletal muscle insulin resistance: A case of imbalanced bioenergetics. Biochim Biophys Acta 2016;1857: 1678–1693.

58. Lundberg JO, Gladwin MT, Ahluwalia A, Benjamin N, Bryan NS, Butler A, et al. Nitrate and nitrite in biology, nutrition and therapeutics. Nat Chem Biol 2009;5: 865–869.

59. Khatri J, Mills CE, Maskell P, Odongerel C, Webb AJ. It is Rocket Science - Why dietary nitrate is hard to beet! Part I: Twists and turns in the realisation of the nitrate-nitrite-NO pathway. Br J Clin Pharmacol 2017;83: 129–139.

